# A compilation of fecal microbiome shotgun metagenomics from hospitalized patients undergoing hematopoietic cell transplantation

**DOI:** 10.1101/2021.08.23.457365

**Authors:** Jinyuan Yan, Chen Liao, Bradford P. Taylor, Emily Fontana, Luigi A. Amoretti, Roberta J. Wright, Anqi Dai, Nicholas Waters, Jonathan U. Peled, Ying Taur, Miguel-Angel Perales, Benjamin A. Siranosian, Ami S. Bhatt, Marcel R.M. van den Brink, Eric G. Pamer, Jonas Schluter, Joao B. Xavier

## Abstract

Hospitalized patients receiving hematopoietic cell transplants provide a unique opportunity to study how the human gut microbiome changes in response to perturbations, and how the resulting changes in the microbiome feedback on its living host. We previously compiled a large-scale longitudinal dataset of stool microbiome compositions from these patients and associated metadata^1^. In that dataset the microbiome analysis was limited to the taxonomic composition of the bacterial population obtained from 16S rRNA gene sequencing. Here, we augment those data with shotgun metagenomic sequences from a nested subset of 395 stool samples. We provide accession numbers that link each sample to the paired-end sequencing files deposited in a public repository, which can be directly accessed by the online services of PATRIC^2^ to be analyzed without the users having to download or transfer the files. We provide examples that show how shotgun sequencing enriches microbiome analyses beyond the taxonomic composition such as the analysis of gene functions including virulence factors and antibiotic resistances, and the assembly of genomes from metagenomic data.

## Background & Summary

The composition of gut microbiome changes in response to mild perturbations such as changes in diet^3^ and strong perturbations such as chemotherapy ^4^ and antibiotics ^5^ that can deplete the majority of the microbes and impact microbiome function^5^. Over the past decades, the microbiome field has sought to characterize compositional changes to perturbations and understand how those changes impact human health^6^. Cross-sectional or longitudinal multi-omics data yielded valuable insights into the population dynamics of gut microbes, their ecological interactions and metabolic functions, and the molecular mechanisms of host-microbe crosstalk^7,8^. Data from patients hospitalized to receive allogeneic hematopoietic cell transplantation (HCT) provide a unique chance to study the gut microbiome in extremely perturbed conditions^9–11^. These perturbations caused by the treatment occur in a planned, scheduled fashion as patients stay in the hospital for several weeks, which enables collecting samples and clinical metadata. The patients receive many drugs including antibiotics that impact the composition and function of the gut microbiome^12,13^. The data also allow us to study how the microbiome composition feeds back on the state of its living host, and address some basic science questions such as how the microbiome influences the dynamics of the human immune system^14^.

We previously published the first data descriptor of our institutional microbiome dataset of HCT patients (> 10,000 samples from >1,000 patients), where we compiled patients’ gut microbiota compositions based on 16S rRNA gene sequencing of fecal samples and its associated metadata^1^. Subsets of this comprehensive dataset were analyzed in a number of publications^9,14–23^. Metagenomic shotgun sequencing is more expensive but has advantages compared to 16S rRNA gene sequencing^24^: it not only reveals the composition of the gut microbiome but also the functions encoded by the genes in the microbiome^25,26^. Bioinformatic tools that analyze shotgun sequencing data for different purposes—taxonomic classification of microbial composition^27^, gene abundance prediction of specialty genes such as antibiotic resistance^28,29^ and virulence factors^29^, genome identification of strain-level or species-level metagenome-assembled genomes (MAGs)^30,31^ and metabolic model reconstruction that translate the DNA sequences to biochemical reactions^32–34^—are now readily available. Some of these tools even work directly with the accession numbers of the sequencing data deposited in public repositories, which greatly facilitates analysis.

Here we compile 395 human fecal samples that were analyzed by metagenomic shotgun sequencing, which is a nested subset of samples we compiled previously and analyzed by 16S rRNA amplicon sequencing^1^. We present examples of functional analyses, including taxonomic composition, gene functions such as virulence factors and antibiotic resistance and the assembly of genomes from metagenomic data. We first conduct a data validation where we check the data for quality by addressing specific questions: Do the compositions inferred from metagenomic and 16S sequencing data agree? How well does metagenomic sequencing capture antibiotic resistance genes? Can the metagenomic data recapitulate the genomic difference of bacterial pathogens? We display the 395 shotgun samples on a t-SNE map of the >10,000 samples of 16S amplicon sequencing^1^. We then investigate correlations between the consistency of stool samples and the read counts of shotgun samples, and we check the correlation of composition between 16S amplicon sequencing and shotgun metagenomes. We then validate the ability to detect antibiotic resistance genes using an orthogonal detection of the *vanA* gene for vancomycin resistance using a PCR test. We used the available tools from PATRIC, a publicly accessible database and tool repository for bacterial genome analysis, to do compositional analysis (kranken2), virulence gene (VFDB) and antibiotic resistant gene (CARD) identification. We assembled metagenomically assembled genomes (MAGs) from shotgun reads and compared them with genomes sequenced from isolates of *Enterococcus faecium* obtained from the same samples^35^. We provide Matlab code to compile the output of these metagenomic analysis tools in a Github repository https://github.com/joaobxavier/shotgun_scientific_data.

## Methods

### Library preparation, shotgun sequencing and human genome decontamination

We compiled 395 of the >10,000 stool samples acquired from allo-HCT patients^1^, extracted the genomic DNA and sequenced on the Illumina HiSeq platform as described previously^14,16^. We removed normal optical duplicates in paired FASTQ files using the clumpify.sh tool from the BBMap package (BBMap – Bushnell B. – https://www.sourceforge.net/projects/bbmap/), producing a pair of read files without duplicates. Using the bbduk.sh script in the BBMap package, we trimmed the right and left side of a read in a pair to Q10 using the Phred algorithm. A pair of reads was dropped if any one of them had a length shorter than 51 nucleotides after trimming. We trimmed 3’-end adapters using a kmer of length 31, and a shorter kmer of 9 at the other end of the read. One mismatch was allowed in this process, and we allowed adapter trimming based on pair overlap detection (which does not require known adapter sequences) using the ‘tbo’ parameter. We used the ‘tpe’ parameter to trim the pair of reads to the same length. We removed human contamination using Kneaddata employing BMTagger. The BMTagger database was built with human genome assembly GRCh38. The paired end read files were uploaded into the Short Read Archive (SRA) of the National Center for Biotechnology Information (NCBI).

### Taxonomy classification and specific gene mapping for metagenomic reads

We used the services provided by the Pathosystems Resource Integration Center (PATRIC)^2^. PATRIC can take input as the SRA accession number of each sample and output the microbiome composition in taxa, as well as genes encoding virulence factors and antibiotic resistances. It uses the algorithm Kraken 2^27^ for taxonomic classification, and the algorithm KMA^36^ to align the metagenomic reads to non-redundant databases. The virulence factor composition analysis is based on the Virulence Factor Database^29^ and the antibiotic resistance composition is based on the Comprehensive Antibiotic Resistance Database (CARD)^28^. The taxonomy, virulence factor and antibiotic resistance table for each of the 395 samples are provided as text tables.

### Genome assembly

We adapted a recently published pipeline to assemble the genomes of bacteria from shotgun sequenced samples^37^. Briefly, the pipeline first assembled contigs using metaSPAdes^38^. Then, it binned the contigs into MAGsusing three different methods: Metabat2^30^ CONCOCT^31^ and Maxbin2 ^39^. The results were then aggregated using DASTool which implements a dereplication, aggregation and scoring strategy^40^ to produce the strain-level genomes.

## Data Records

The shotgun sequenced samples were deposited in the NCBI/SRA as paired-end fastq files decontaminated of human reads. We updated the data table tblASVsamples.csv in Figshare (https://doi.org/10.6084/m9.figshare.12016983.v8) that we had previously published as part of our microbiota compilation^1^: We added a new column to the table, ‘AccessionShotgun’, which lists the SRA accession record for each of the 395 samples presented here. All other samples were left with an empty entry in column ‘AccessionShotgun’. The table can be updated in the future as new shotgun sequences become available.

We compiled the additional tables for each sample as comma-separated value (csv) files in Figshare (https://figshare.com/account/home#/projects/120102) as following:

- ReadCounts.csv: list the 395 samples used this study for shotgun
  - SampleID: Name of samples
  - Readcount: Number of reads for each sample after decontamination of human reads.
- Abundance: A Kraken 2 report provides information of the bacterial taxa in each sample.
  - Kindom, Phylum, Class, Order, Family, Genus: Each column contains name of taxonomic classification of each sample
  - ColorOrder: Numeric data representing the order that each taxa was plotted in our manuscript.
  - HexColor: Color code in hex format for each taxa
  - 395 columns using sample names as column names: Numeric data as the relative abundance of each sample. Every column should sum to 1.
    - NOTE: The (U)nclassified reads from kraken2 are not included in the calculation.
    - The sample names of those columns are modified to be compatible with the format requirement in matlab. To convert the names back to match the SampleID in the rest of the files, ‘s’ at the beginning of each column name should be removed, and underscore (’_’) needs to be converted to period (eg. ‘sFMT_0001A’ will become ‘FMT.0001A’).
- CARD.csv: A table provides information of the antibiotic resistance genes in each sample.
  - Template: Hit of resistant genes in CARD
  - Accession: NCBI accession number of the template
  - Genome: Strain names where the template gene is found
  - Species: The species of the strain
  - resistGene: Gene name if available.
  - resistMechanism: Mechanism of resistance interpreted by CARD
  - Zoliflodacin-unknown (multiple columns): Antibiotics whether the gene is predicted to be resistant to, including unknown.
  - Score, Expected, Template_length, Template_Identity, Template_Coverage, Query_Identity, Query_Coverage, Depth, q_value, p_value: Parameters reported by CARD to show how well the match is.
  - shotgunReadcount: Number of reads for each sample after decontamination of human reads.
  - RelavantPercentInCARD: The number of reads matched to the template resistant gene / Total reads matched to all CARD genes in the sample
  - PercentageInShotgun: The number of reads matched to the template resistant gene / Total reads in the sample
  - Mutation: Information whether the antibiotic resistance is conferred by mutation
  - SampleID: Sample ID for each shotgun sequencing
- VFDB.csv: A VFDB report provides information of the virulence factors in each sample.
  - Template: Hit of virulence genes in VFDB
  - Function: Predicted function of the template
  - Genome: Strain names where the template is found
  - Score, Expected, Template_length, Template_Identity, Template_Coverage, Query_Identity, Query_Coverage, Depth, q_value, p_value: Parameters output by VFDB that report how well the match is.
  - shotgunReadcount: Number of reads for each sample after decontamination of human reads
  - RelavantPercentInVF: The number of reads matched to the virulence gene / Total reads matched to virulence genes in the sample
  - PercentageInShotgun: The number of reads matched to the virulence gene / Total reads in the sample
  - SampleID: Sample ID for each shotgun sequencing
- Taxa_Not_In_16S.csv: Taxa absent in 16S gene sequencing but present in shotgun metagenomic sequencing in our 395 samples
  - TaxaName: Name of missing taxa
  - Taxa: Classification level of taxa names (eg. Genus)
  - frequencyPresentInShotgun: Counts of non-zero abundance found in shotgun sequencing(non-zero) / Total number of samples
  - medianRelAbdInShotgun: Median of relative frequency in shotgun metagenomic sequencing of that taxa
- Taxa_Not_In_Shotgun.csv: Taxa absent in shotgun sequencing but present in 16S gene sequencing in our 395 samples
  - TaxaName: Name of missing taxa
  - Taxa: Classification level of taxa names (eg. Genus)
  - frequencyPresentIn16S: Counts of non-zero abundance found in 16S (non-zero) / total number of samples
  - medianRelAbdIn16S: Median of relative frequency in 16S sequencing of that taxa

## Technical Validation

### The nested subset of shotgun-sequenced samples explores various microbiome states experienced by patients

Out of the >10,000 samples with 16S rRNA gene sequencing^1^, a total of 395 samples were sequenced using metagenomic shotgun technique for the purposes of different projects. Using a t-SNE map generated from Bray-Curtris dissimilarity matrix of 16S rRNA gene sequencing (**Fig. 1a**), we highlighted the samples with shotgun sequencing data available, which are distributed across the map (**Fig. 1b**). The nested subset captures a wide range of microbiome states, representing many states found in the original dataset of >10,000 samples. For example, both *Enterococcus*-dominated “dysbiotic” states (dark green portion of t-SNE projection in **Fig 1a**) as well as the “healthier” Clostridia-enriched states (grey portion of t-SNE projection in **Fig 1a**) are well-represented by the nested shotgun dataset.

**Figure 1.**
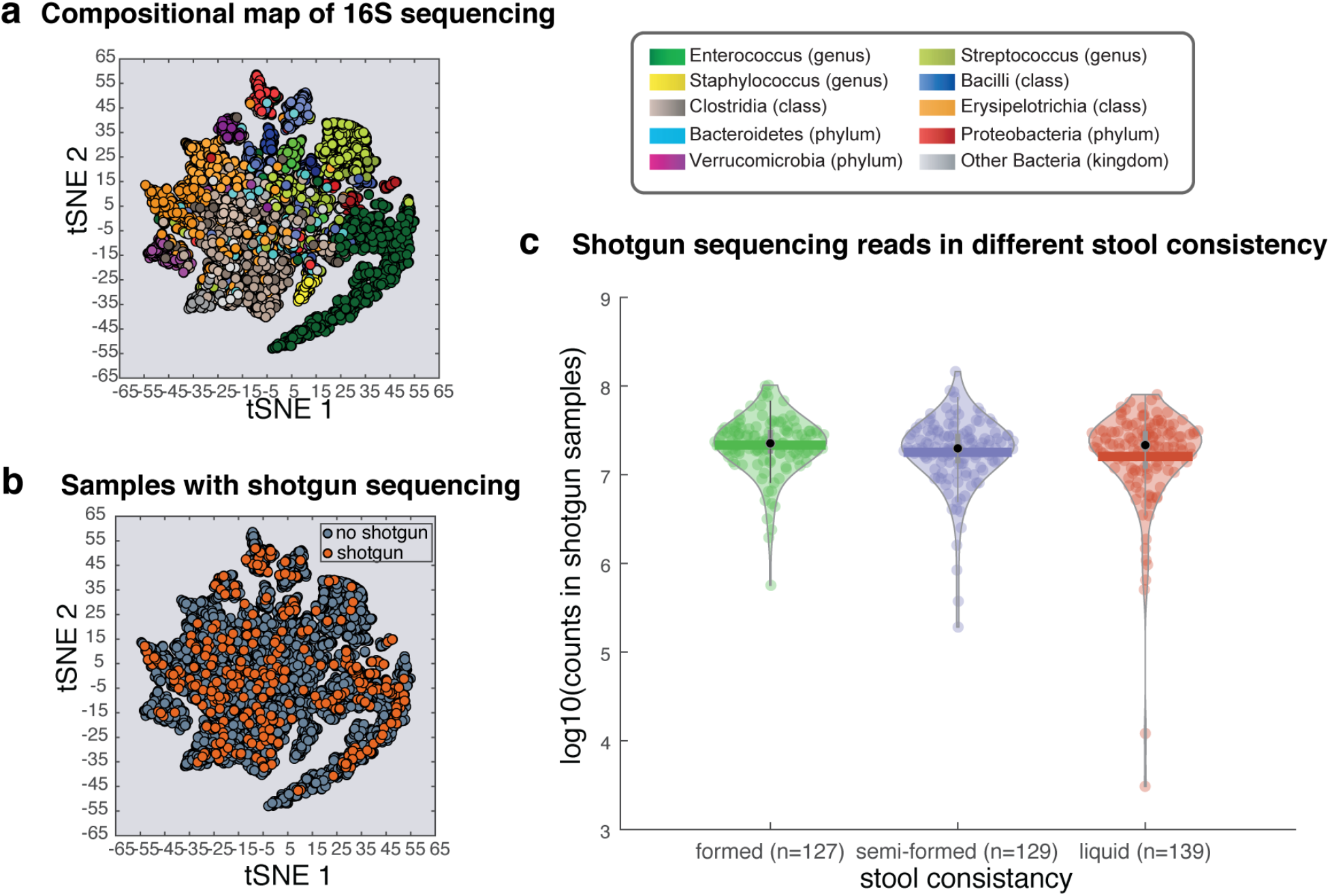
The metagenomic samples cover the majority of microbiome compositional states observed in fecal samples from allo-HCT patients. **a.** The t-SNE plot built using the taxonomic composition obtained by 16S amplicon sequencing of N samples from N unique patients^1^; the different colors indicate the most abundant taxon in each sample. **b.** Location of nested subset of N samples from N unique patients with shotgun sequencing is broadly distributed across the entire map. **c..** The sequencing depth of shotgun sequenced samples varies between 10^6^ reads to 10^8^ reads, with outliers in the liquid samples whose microbiome may yield different sizes of libraries.

The stool consistency from these patients varies widely. At the time of stool aliquoting, stool consistency was assessed by laboratory technicians using a scale of “formed, semi-formed, and liquid” to indicate the dry weight of stool^41^. The link between stool consistency and gut microbiota composition has been examined in the 16S amplicon sequencing pipeline ^42^. Before diving into the diversity analysis, we first tested if the stool consistency would associate with the read count of shotgun sequencing. We observed that the median of the three types of stool (formed, semi-formed and liquid) are all above 10^7^ reads per sample (**Fig. 1c**), except for two samples from the liquid group that showed lower reads count than the rest of the samples (<10^5^).

### Validating the taxonomic composition of the shotgun metagenomes

We first sought to compare the taxonomic classifications obtained by 16S rRNA gene sequencing^1^ and shotgun sequencing analyzed by Kraken 2. A visual inspection of the bacterial compositions suggests that the two ways to analyze the taxonomic composition of the bacterial population agree well: When we compare the compositions from the patient with the highest number of collected samples we can see a reasonable match (**Fig. 2 a,b**) between stacked bar plots of compositions color-coded according to a palette designed to highlight microbiome injury patterns^1^.

**Figure 2.**
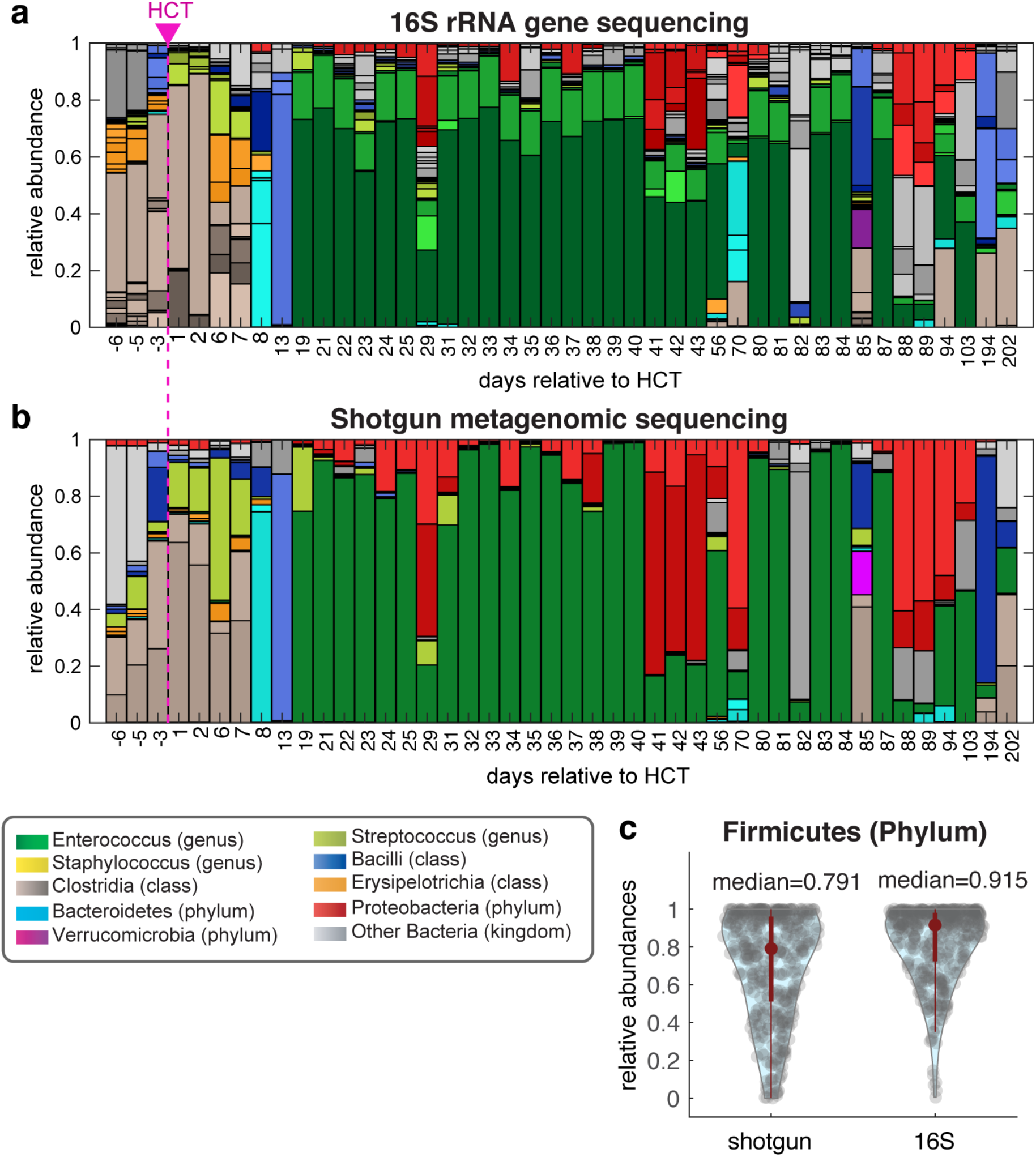
Taxonomic composition of the microbiome in patient stool samples agrees in general between shotgun sequencing and 16S rRNA amplicon sequencing, with some notable differences. **a,b.** The taxonomic composition is determined by 16S rRNA sequencing (A) and shotgun metagenomics (B) for the samples from a single patient (PatientID 1252). The samples are ordered in time and the dashed line separates the samples collected before and after allo-HCT. **c.** The median composition (red dot) in Firmicutes can be notably different when determined using the two approaches (ranksum test, p<0.05).

A closer inspection shows however that the Shotgun sequencing missed some of the taxa seen in the shotgun data (eg. the orange bar representing *Erysipelotrichia* in day 56, and the blue bar representing *Bacilli* in day 82). We then compared the relative abundance of different taxa as assessed by 16S and shotgun sequencing and identified the taxa with median relative abundance higher than 10% and significantly different between 16S and shotgun sequencing, among which the Firmicutes (phylum) has the overall highest abundances (**Fig. 2c**), which could be explained by its higher copy number of rRNA in the genome ^43^. The other taxonomic groups are Bacilli (class), Clostridia (class), Clostridiales (order) and Lactobacillales (order).

There were some taxa only found in either approach, and the shotgun sequencing found much more taxa than the 16S gene sequencing (1870 missing in 16S but present in shotgun; 182 missing in shotgun but present in 16S; the .csv files can be found in Figshare). There are a few reasons that could possibly explain the disagreement between 16S and metagenomic shotgun sequencing: First is the ambiguous naming where a taxa was renamed later (e.g. Enterobacteriales was renamed to Enterobacterales), sharing the same sequencing of tested 16S region (Escherichia and Shigella are the same in 16S gene sequencing), and poorly studied taxa (e.g. ‘CAG-352’ in 16S sequencing). The second is the detection variation. The taxa missing entirely either in 16S or in shotgun overall have very low median abundance even detected in the other pipeline – only ‘Incertae Sedis’ and ‘CAG-352’ found in 16S are between 1 to 4 percent, while all other missing taxa show less than 1% median relative abundance in the other pipeline where they are found. The third reason could be differences between the databases used for taxonomy detection. To systematically compare 16S and shotgun sequencing, we calculated correlation of relative abundance between the two pipelines in taxonomic classification. The agreement between 16S and metagenomic shotgun were generally high and decreased for lower taxonomic ranks (**Fig. 3**), indicating that the two approaches have different sensitivities for taxonomy. The few samples with low correlation tended to be those sequenced at a lower read depth, but some samples sequenced deeply could also have low correlation (**Fig. 3a-e**), indicating that other factors than read counts may affect the taxonomic mapping of metagenomes.

**Figure 3.**
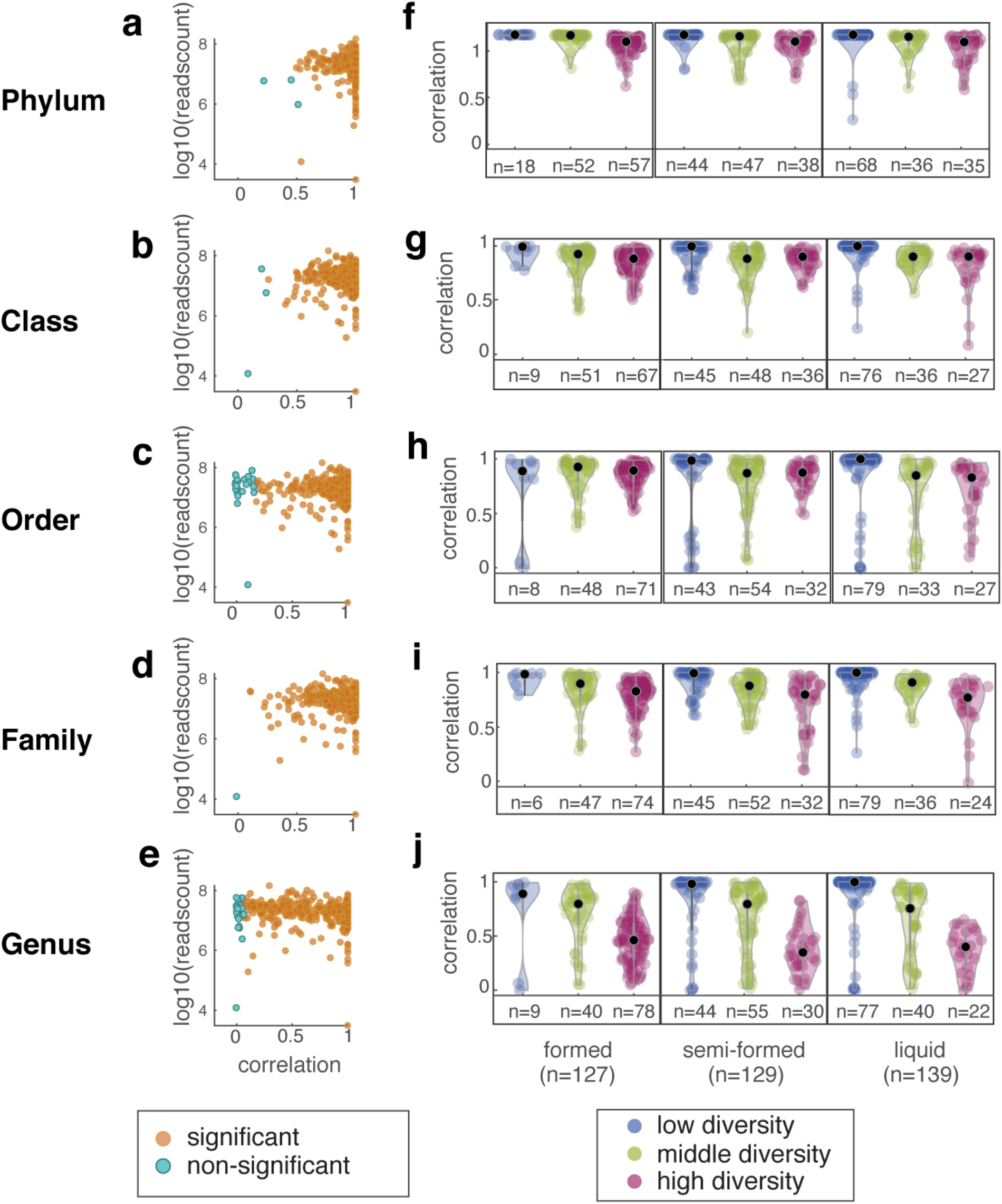
Correlation between the taxonomic classifications obtained by shotgun sequencing and 16S rRNA amplicon sequencing. The correlation between the two approaches is different at each taxonomic level, but seems unaffected by the read depth of each sample (**a-e**). Formed stool mainly contains samples with higher diversity, and high diversity samples usually display lower correlation between the two sequencing pipelines (**f-j**). Each point is a taxon from one of 395 samples. Black dots in f-j indicate the median of each category. The numbers on the x-axes display the number of samples in different diversity groups.

One possible explanation is that discrepancies between the database of bioinformatic pipelines may become especially visible for highly diverse samples, leading to low correlations. We therefore calculated the alpha-diversity as determined by the Shannon index for each taxonomic classification. Then we divided the values into three groups: high diversity (top 33%), middle diversity and low diversity (bottom 33%). Because stool consistency is a marker of species richness in microbiome^42^, we stratified our samples by stool consistency and examined if the diversity clusters are discrete among different consistencies (**Fig. 3f-j**). The high diversity group does have overall lower correlation, which becomes more and more obvious from phylum to genus. We also noted 5 samples with both low diversity (in bottom 33% percentile) and low correlation (less than 0.01) between the taxonomies quantified by shotgun and 16S. Four samples with the lowest correlation (<0.001) in genus composition are also in the bottom 33% percentile of diversity, which were caused by a failure by the 16S pipeline to detect a bacterium of the genus *Klebsiella*. This example illustrates a possible source of error in the 16S pipeline that may be improved using shotgun metagenomics.

### Validating the detection of antibiotic resistance genes using a PCR to detect the *vanA* gene

PATRIC provides web services to quantify virulence factors and antibiotic resistance genes in the microbiome samples. To test how well this analysis detected antibiotic resistant genes, we used PCR to detect the presence of an important gene for vancomycin resistance *vanA^1^* (**Fig. 4a**). Vancomycin is a glycopeptide antibiotic that is given to many of the allo-HCT patients in this cohort as prophylaxis to prevent infections by *Streptococcus*^44^. The samples chosen for the PCR test contain ≥2% enterococcal sequences, since the presence of vanA is usually a sign of *Enterococcus* domination^45,46^. We compared the relative abundance of the *vanA* gene, as quantified by the PATRIC analysis of CARD genes, in *vanA* PCR(−) versus *vanA* PCR(+) samples and we saw a significant agreement (**Fig. 4b**). In the PCR(+) group only 2 samples (out of 120) have zero abundance of *vanA* in metagenomes while 143 out of 186 are zero in PCR(−) group, suggesting that shotgun sequencing may be more sensitive than the PCR.

**Figure 4.**
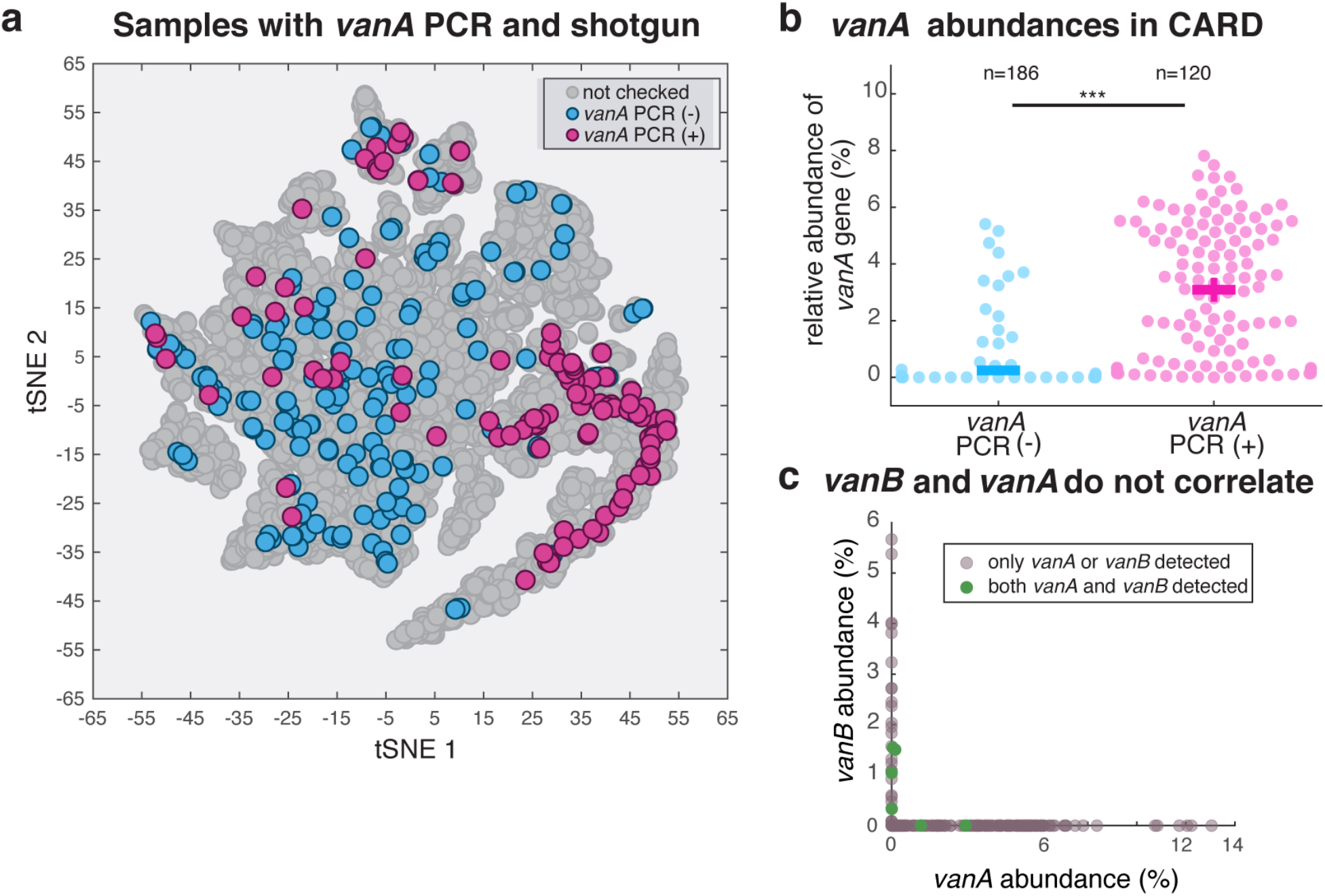
The shotgun sequencing detects the presence of antibiotic resistance genes, using the PATRIC service with the CARD database. **a.** Localization of the *vanA*(+/−) samples in 16S clustering map shows a high concentration of *vanA*(+) samples in the region of domination by *Enterococcus* (green in Fig.1a). **b.** PCR(+) samples have higher relative abundance of the *vanA* gene detected by shotgun sequencing. **c.** The *vanA* and *vanB* genes are practically mutually exclusive in patients’ stool samples. The samples with two genes simultaneously detected represent a very small fraction of the total samples. The abundances of the two genes are not correlated.

There are other genes besides *vanA* important for resistance to vancomycin. We examined whether the abundance of another vancomycin resistance gene, *vanB*, correlated with *vanA*. We saw that although those two genes are negatively correlated in our gut microbiome samples (r=-0.28, p<0.05), plotting the gene abundance (**Fig. 4c**) reveals that only six samples carry both *vanA* and *vanB* (**Fig. 4c**, green dots). In all the other cases, only *vanA* or *vanB* was found, suggesting that bacteria harboring these genes may be excluding invasion by competitors harboring the other gene.

### Validating the assembly of genomes from shotgun sequences

New bioinformatic pipelines have enabled us to assemble the genomes of bacteria from shotgun sequences^37,47,48^. To illustrate the utility of our data for this type of analysis, we ran a published metagenomic analysis pipeline to find MAGs (metagenome-assembled genomes) from the samples of PatientID 1044, where we know from genotyped isolates obtained in a previous study that the patient carried *Enterococcus faecium* in the gut^35^. We found 7 high-quality MAGs classified as *E. faecium*, each from a different stool sample and has a completeness higher than 95%. We then compared these MAGs with the genomes of our 26 *E. faecium* isolates^35^ in a phylogenetic tree (**Fig. 5**). Our previous study^35^ had shown that the patient contained at least two distinct strains of *E. faecium*. The MAGs confirmed the observation: Three MAGS, MAG_1044M_maxbin0003, MAG_1044P_maxbin002 and MAG_1044L_4, located in the tree branch of the strains from the same three samples that collected in the later days relevant to HCT, whereas four MAGs, MAG_1044J_16, MAG_1044G_10, MAG_1044H_27 and MAG_1044I_maxbin001, located in the tree branch of the other strains that were isolated from the same four samples from the earlier days after HCT. The comparative analysis indicates that the dominant *E. faecium* strain has shifted between day 5 and day 7 after HCT.

**Figure 5.**
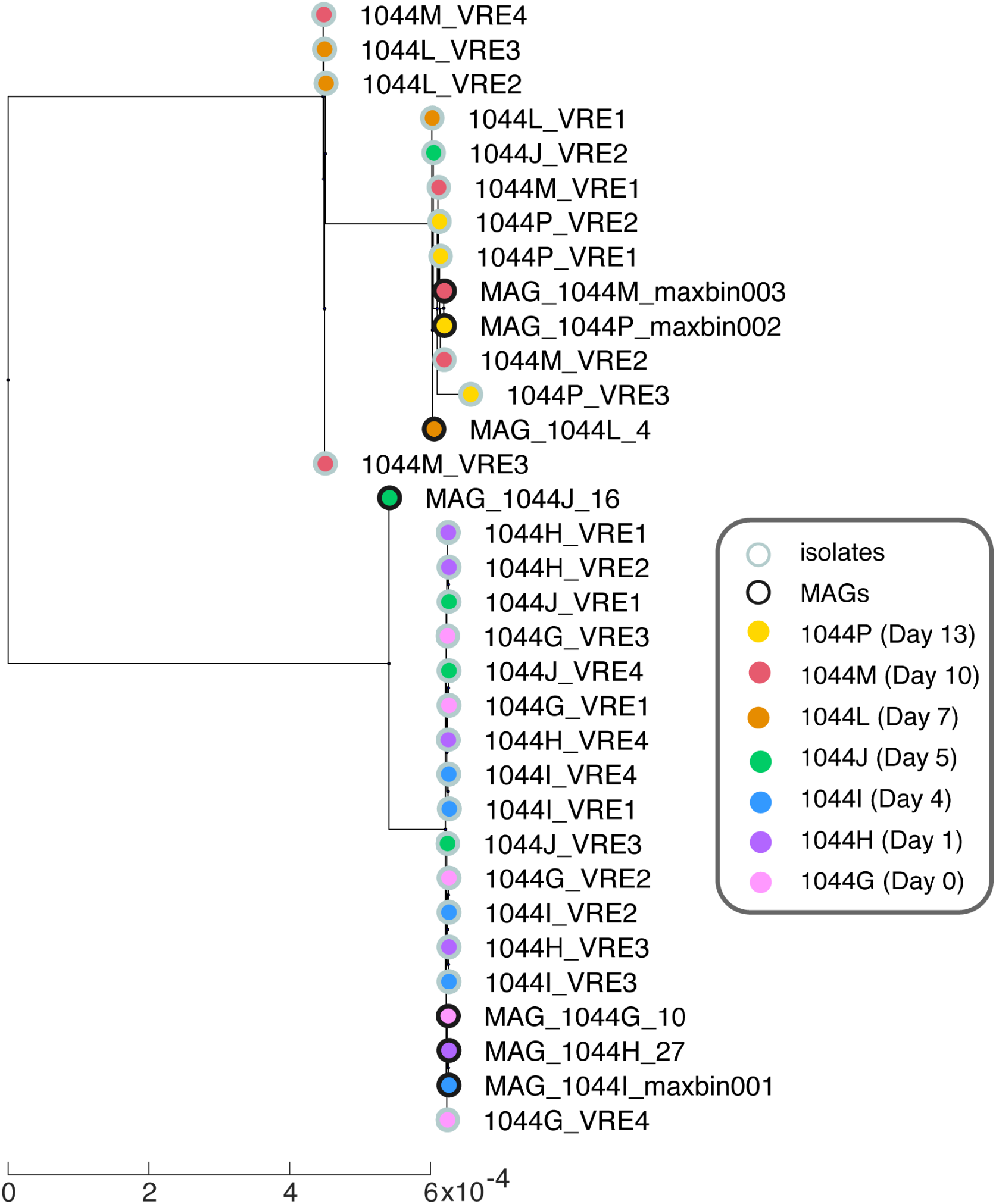
Shotgun sequencing data provide metagenomically-assembled genomes (MAGs) that compare well with the genomes of isolates from the same patient stool samples. MAGs from *E. faecium* obtained from different samples collected from patient 1044 reveal an intraspecies diversity. The phylogenetic tree contains the 7 MAGs and 26 *E. faecium* genomes obtained from isolates and analyzed in a previous study^35^. The number of days after each sample is the day relative to the HCT of this patient.

## Code Availability

The customized analysis code (Matlab 2020a) used for the examples provided below is available in the GitHub repository (https://github.com/joaobxavier/shotgun_scientific_data). with each part in a separate directory:

- Figure 1: Figuer1/scFigure1.m
- Figure 2: Figuer2/scFigure2.m
- Figure 3: Figuer3/scFigure3.m
- Figure 4: Figuer4/scFigure4.m
- Figures 5: Figuer5/scFigure5.m

**Acknowledgements, Author Contributions & Competing Interests**

## Acknowledgements

This work was supported by the National Institutes of Health (NIH) grants U01 AI124275 to EP and JBX and R01 AI137269 to JBX and YT. This work could not have been possible without our longtime collaborator Eric Littman. Eric passed away in June 2021 when we were starting to prepare this manuscript and he is not on the author list only because of the need to have all authors read and agree on the final version. We acknowledge Eric’s invaluable contribution to this dataset and his analyses of shotgun sequencing data that inspired some of the examples presented here.

## Author contributions

J.Y.,C.L. and J.B.X. wrote the manuscript. J.Y., C.L., and J.B.X. designed the analyses. Y.T. contributed to the clinical data preparation, B.P.T. provided the 16S data processing pipelines and did some of the preliminary analysis that led to this work. E.F., L.A.A. and R.J.W. processed patients’ stool samples. All authors contributed to the writing and interpretation of the results.

